# Unveiling the mechanisms of black phosphorus nanosheets-induced viable but non-culturable state in *Bacillus tropicus*

**DOI:** 10.1101/2024.06.17.599389

**Authors:** Zhiqiang Xiong, Qing Zhao, Ming Zhao, Liwei Liu, Jin Zeng, Siyu Zhang, Shuo Deng, Daxu Liu, Xuejiao Zhang, Baoshan Xing

## Abstract

The release of black phosphorus (BP) nanosheets has raised concerns regarding potential ecological risks. Previous studies have confirmed their toxicity to bacteria, but discrepancies were observed between results obtained from the growth curve and colony forming unit (CFU) methods, indicating the possibility of bacterial cells entering a viable but non-culturable (VBNC) state induced by BP nanosheets. To accurately assess the risks, it is crucial to understand the underlying mechanisms. In this study, we investigated the effect of BP nanosheets on *Bacillus tropicus*, a gram-positive bacterium, using transcriptome sequencing and biological assays. Our findings revealed that BP nanosheets caused minimal cell death but predominately induced the VBNC state in most cells. At the transcriptional level, we observed significant down-regulation of pathways associated with cellular metabolism and respiratory chain in response to BP nanosheet treatment. Bacterial cells in the VBNC state exhibited depressed respiration to maintain basal cellular activity. Additionally, the reduced cellular respiration and metabolic activity were associated with a decrease in antibiotic susceptibility of the bacteria. These results provide new insights into the antibacterial mechanisms of BP nanosheets and emphasize the necessity of employing appropriate approaches, beyond the traditional CFU method, to assess the bacterial toxicity of nanomaterials.

**Environmental implication:** Bacteria play a crucial role as indicators in ecological risk assessment. Although numerous studies have highlighted the exceptional antibacterial properties of black phosphorus (BP) nanosheets, the unique viable but non-culturable (VBNC) state of bacteria is often overlooked when evaluating the ecological risks of nanomaterial, including BP nanosheets. In our study, we found that BP nanosheets can induce *Bacillus tropicus* into a VBNC state by suppressing cellular metabolism- and respiratory chain-related pathways, shedding light on their ecological risk assessment implications. This finding underscores the importance of utilizing appropriate approaches in evaluating the bacterial toxicity of nanomaterials.

## 1. Introduction

Black phosphorus (BP) nanosheets, an emerging type of two-dimensional (2D) nanomaterial, have gained wide attention due to their excellent optical and electrical properties, as well as their favorable biocompatibility. These features make them highly promising for a wide range of applications, including optoelectronic devices,^1, 2^ photocatalysis,^3^ biomedicine,^4^ and antibacterial materials.^5^ Advances in manufacturing technology have enabled the cost-effective production of BP-based nanomaterials on a gram scale,^6, 7^ laying the foundation for their large-scale utilization. However, the increasing production and consumption of BP nanosheets raise concerns about their potential release into the environmental. Bacteria, as vital components of the ecological environment, play a crucial role in the field of ecological nanotoxicology.^8^ Therefore, investigating the bacterial toxicity of BP nanosheets is not only essential for the assessment of their ecological risk, but also provides indispensable insights for the development of BP-based antibacterial materials.

Previous studies have commonly used the colony forming unit (CFU) method to evaluate the bacterial toxicity of nanomaterials, including BP nanosheets, due to its simplicity and directness.^9–11^ However, this method is limited to detecting cells under cultivable condition. Bacterial cells that do not grow on culture media are usually considered dead or irreversibly inactivated in nanotoxicity research.^12^ Nevertheless, it is recognized that non-growing bacteria may still retain metabolic capacity despite their inability to form individual colonies on standard laboratory media. This state is referred to as the viable but non-culturable (VBNC) state.^13^

Bacterial cells have the ability to enter a VBNC state as a survival strategy in response to adverse environmental stresses, such as unfavorable temperature,^14^ pH,^15^ or exposure to nanomaterials.^16^ Transcriptome sequencing has been widely employed to elucidate the evolutionary mechanisms of stress-induced VBNC cells.^17^ For instance, Zhao et al. discovered that *Escherichia coli* (*E. coli*) cells decreased adenosine triphosphate (ATP) generation by altering pyruvate catabolism when transitioning into VBNC state under high-pressure of CO_2_. In response, the bacterial cells elevated the degradation of L-serine and L-threonine, leading to increased adenosine monophosphate (AMP) generation.^18^ In addition, the initiation of the VBNC state can also be activated by stimulus-response mechanisms in the presence of certain chemicals.^17^ Bacteria sense external stress through membrane-bound histidine-kinases, with response regulators orchestrating cellular responses via modulating the expression of specific target genes.^19^ *E. coli* cannot enter the VBNC state under osmolarity, pH, and starvation stresses without the expression of histidine kinases.^20^ Furthermore, the induction mechanisms of the VBNC state have been elucidated in various bacterial strains, including *E. coli*, *Lactobacillus acetotolerans*, and *Rhodococcus biphenylivorans*, with studies demonstrating that changes in gene expression in VBNC cells depend on the specific inducer.^17^

Several studies have shown that nanomaterials can induce bacteria to enter the VBNC state. For example, silver nanoparticles have been identified as inducers of the VBNC state in *Pseudomonas aeruginosa* (*P. aeruginosa*).^12^ Exposure to silver nanoparticles completely inhibited the culturability of *P. aeruginosa*, while the metabolic activity, total cell counts, and cytoplasmic membrane integrity remained unaffected. Similarly, the presence of VBNC cells of *Staphylococcus aureus* (*S. aureus*) and *P. aeruginosa* has been observed under stress induced by graphene nanoplatelets, as determined by the difference between the number of viable and culturable cells.^21^ Although BP nanosheets have also been extensively studied for their antibacterial effects, most studies have only utilized the CFU method to evaluate their antibacterial activity.^9, 10^ In our previous studies, we noticed a significant discrepancy between the results obtained from the growth curve and CFU methods,^22,23^ indicating that BP nanosheets may induce bacteria to enter the VBNC state. This speculation is supported by similar findings reported by Naskar et al.^11^ However, the underlying mechanisms of this response to BP nanosheets and the induction of the VBNC state by nanomaterials have not been clearly elucidated.

In this study, we investigated the response of *Bacillus tropicus* (*B. tropicus*), a gram-positive bacterium, to BP nanosheets. Through the integration of transcriptome sequencing and complementary biological tests, including bacterial toxicity assessments, cell morphology evaluations, measurement of lactate dehydrogenase (LDH) release levels, cell respiration analysis, and resuscitation assays, we discovered that BP nanosheets have the ability to induce bacteria to enter the VBNC state. Moreover, our findings revealed that this phenomenon was driven by the down-regulation of pathways related to the respiratory chain and cellular metabolism. This study significantly contributes to our understanding of the molecular mechanisms underlying the bacterial toxicity of BP nanosheets and provides valuable insights for assessing the environmental risks associated with their release.

## 2. Experimental section

### 2.1 Antibacterial test

Gram-positive *B. tropicus* was used as a model strain to investigate the bacterial toxicity of BP nanosheets. *B. tropicus* was cultured overnight at 37 °C with a shaking speed of 120 rpm, and the bacterial concentration was determined by the CFU method. The cultured bacterial cells were then harvested and added into 10 mL of fresh Luria-Bertani (LB) medium containing different concentrations of BP nanosheets (1, 10, 50, 100 µg mL^-1^) to reach the final bacterial concentration of 10^6^ CFU mL^-1^. These co-culture mixtures were incubated on a shaking incubator at 120 rpm and 37 °C. At specific time points (2, 4, 6, 8, 10, 12 h), samples were taken from the co-culture mixtures and the optical density (OD) value was measured at a wavelength of 600 nm on a Microplate reader (BioTek, USA) to record the growth curve. Bacterial cells cultured in sterile water without BP nanosheets were set as control. To eliminate absorbance caused by BP nanosheets, control samples containing only different concentrations of BP nanosheets in equal volumes of sterile water without any bacterial cells were also prepared. After 12 hour of incubation, 1 mL of the bacterial suspension was sampled, and 100 µL of 10-fold serial dilutions was spread onto agar plats. The plates were then incubated at 37 °C for 12 h, and the number of CFUs was counted. The bactericidal efficiency (%) was calculated as follows:

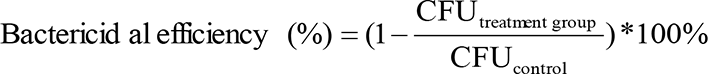

### 2.2 Bacterial viability assay

To assess the viability of *B. tropicus*, the ATP production was measured with the BacTiter-Glo microbial cell viability assay kit. This assay relies on the fluorescence intensity, which is proportional to the amount of ATP present in the cells.^16^ After treatment with different concentrations (0, 1, 10, 50, 100 μg mL^-1^) of BP nanosheets, the bacterial cells were collected by centrifugation at 10000 rpm for 10 min at 4 °C. After wished by sterile PBS for at least 3 times, the bacterial dispersions were diluted 1:100 in sterile PBS. Subsequently, 50 μL of the diluted cells and 50 μL of the ATP kit reagent were added to each well of a 96-well microtiter plate. The plate was then briefly mixed on an orbital shaker and incubated for 5 minutes. Following the incubation, the fluorescence intensity was measured by a multi-function microplate reader (LAI-2000, Tecan, Austria).

### 2.3 Calculation of the percentage of VBNC cells

To calculate the percentage of VBNC cells, all assays were performed with an identical sample set. The total bacterial count in each treatment group, comprising both viable and dead cells, was calculated by multiplying the total bacterial count in control group and the growth activity of the treatment group from the results of growth curves (Figure 1a). The amount of viability cells, which includes both culturable and VBNC cells, was obtained by multiplying the total bacterial count in control group and the bacterial viability of the treatment group from the results of ATP production. The number of culturable cells was determined by multiplying the total bacterial count in control group and the cell viability of the treatment group determined by the CFU method. Consequently, we calculated the amount of VBNC cells by subtracting the number of culturable cells from the total viable count. We also calculated the amount of dead cells by subtracting the number of live cells from the total bacterial count. The percentage of cells in each state was obtained by comparing the cell count in each state to the total cell count for each treatment group. The detail process can be found in the supporting information.

**Figure 1.**
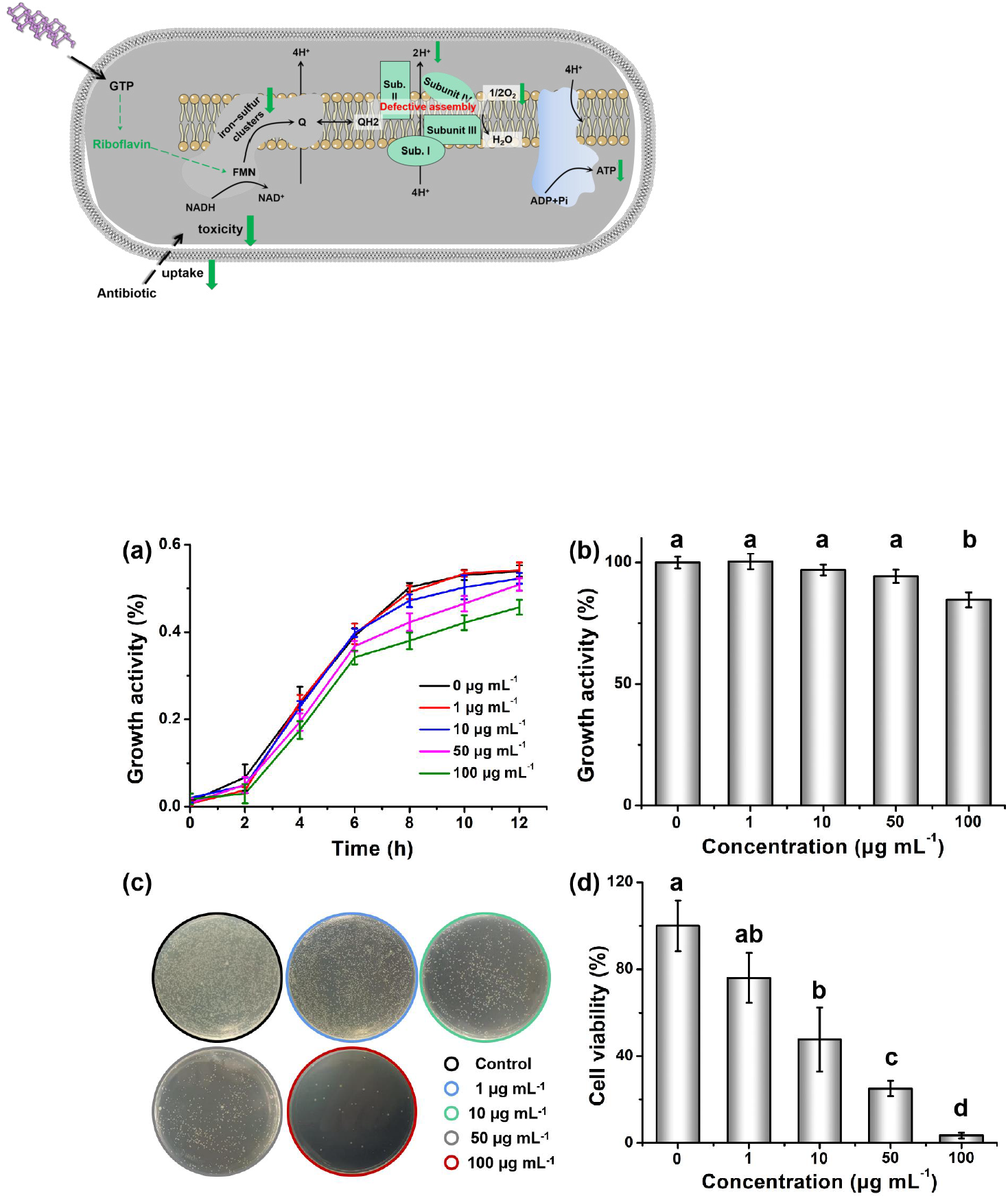
(a) Growth curves under the treatment of different concentrations of BP nanosheets. (b) Growth activity after 12 hours of exposure to BP nanosheets. (c) Images of colonies after 12 hours of exposure to different concentrations of BP nanosheets. (d) Bacterial viability assessed by the CFU method. Images of colonies after 12 hours of exposure to different concentrations of BP nanosheets.

## 3. Results and discussion

### 3.1 Effect of BP nanosheets on bacterial viability

The impact of BP nanosheets on the growth activity of *B. tropicus* was assessed using both growth curve measurements and the CFU method. The results showed that at concentrations of 1 and 10 μg mL^-1^, BP nanosheets had minimal effect on the growth of *B. tropicus*. However, as the concentration increased, the growth activity of *B. tropicus* was moderately reduced. Specifically, exposure to 100 μg mL^-1^ of BP nanosheets for 12 h resulted in a 15.32% ± 3.10% decrease in growth activity compared to the control (Figure 1a,b). In addition, the CFU method was employed to evaluate the impact of BP nansoheets on the culturability of *B. tropicus*. It was observed that the culturability of *B. tropicus* decreased in a concentration-dependent manner, with reductions of 23.95% ± 11.49%, 52.34% ± 14.80%, 74.93% ± 3.56%, and 96.57% ± 1.33% at concentrations of 1, 10, 50, and 100 µg mL^-1^, respectively, after 12 h (Figure 1c,d). These findings aligned with previous studies that demonstrated the antibacterial activity of BP nanosheets against *E.coli* and *Bacillus subtilis* (*B. subtilis*), showing inhibition rates ranging from 91% to 99% at a dose of 100 µg mL^-1^.^9, 23^

However, a notable disparity was observed between the results obtained from the growth curve and the CFU methods, indicating a discrepancy in bacterial activity and culturablity (Figure 1b,d). While the growth curve suggested moderate growth inhibition induced by BP nanosheets, the CFU method showed extremely low values. Moreover, we assessed the survival of bacteria using SYTO/PI staining. The results demonstrated that the majority of *B. tropicus* cells remained alive after co-incubation with BP nanosheets (100 µg mL^-1^) for 12 h (Figure 2a,b, Figure S1). These cells exhibited green fluorescence intensity similar to the control group, with only slightly stronger red fluorescence. The quantification results of the fluorescence intensity indicated that BP nanosheets increased the percentage of dead cells from 4.74% ± 0.10% to 6.84% ± 0.66% compare to control (Figure 2d). In addition, only a fraction of bacterial cells showed compromised membrane integrity after 12 hours of treatment with BP nanosheets (100 μg mL^-1^), while many cells maintained intact cytoarchitecture with smooth membranes, resembling the control cells (Figure 2c).

**Figure 2.**
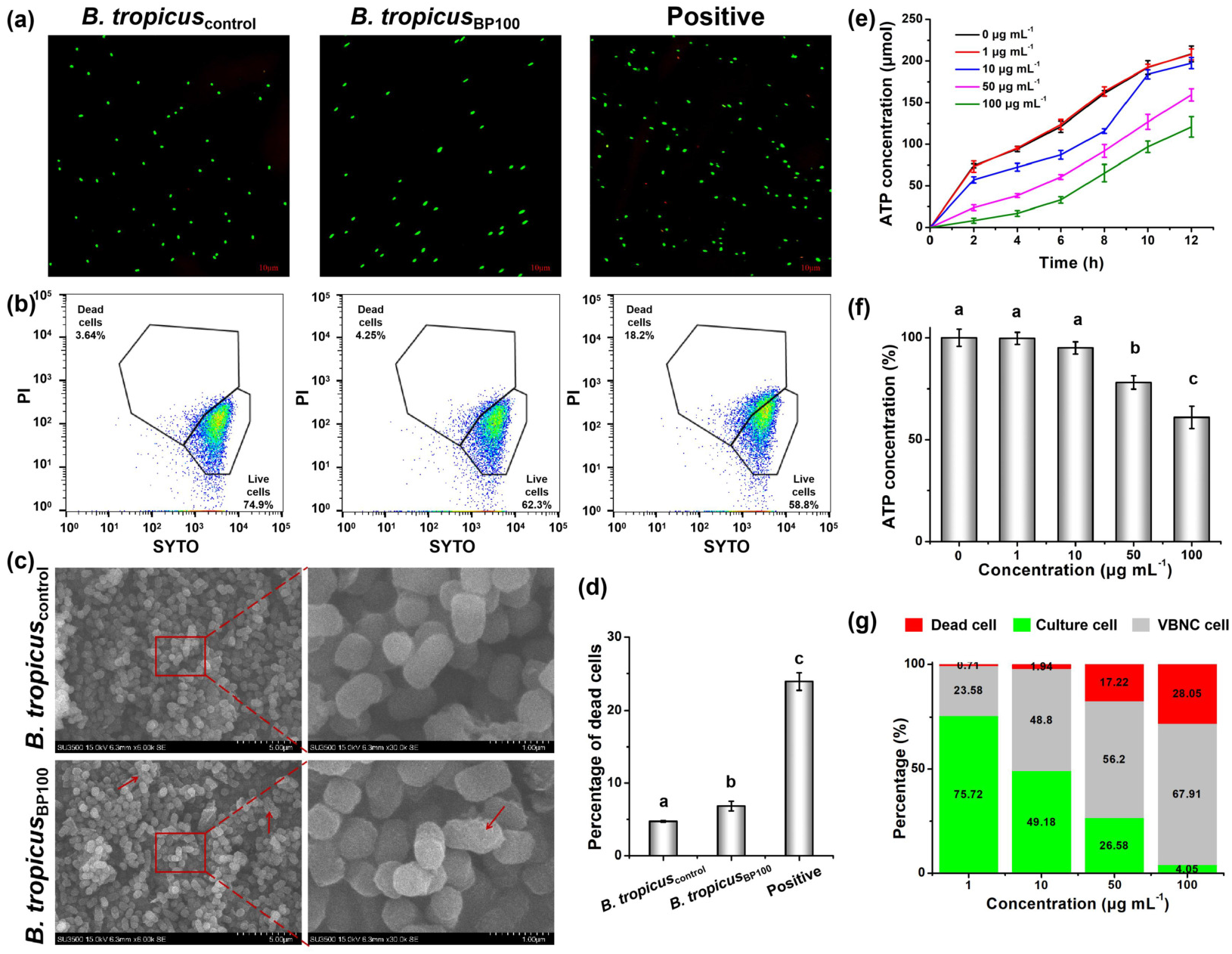
(a) Fluorescence images of *B. tropicus* cells before and after treatment with 100 μg mL^-1^ of BP nanosheets for 12 h, visualized by SYTO/PI staining. *B. tropicus* cells treated with 0.1 H2O2 for 2 h were set as positive control. Flow cytometry analysis of *B. tropicus* cells using SYTO/PI staining (b) and the PI fluorescence intensity in *B. tropicus* cells (d) after treatment with 0 and 100 μg mL^-1^ of BP nanosheets and 0.1% H2O2 for 12 h. (c) Scanning electron microscopy (SEM) images of *B. tropicus* before (upper) and after (under) incubation with 100 μg mL^-1^ of BP nanosheets for 12 h. Scale bars in the images and enlarged views were 5 and 1 μm, respectively. (e) Growth curves determined by measuring the ATP concentrations following the treatment with different concentrations of BP nanosheets. Bacterial viability based on ATP detection (f) and percentage of *B.tropicus* cells in different states (g) after 12 hours of exposure to BP nanosheets.

Based on these findings, it is plausible to consider the possibility of *B. tropicus* entering a specialized bacterial state known as the VBNC state. In this state, bacteria exhibit very low metabolic activity, rendering them unable to form colonies.^16^ The VBNC state is regarded as a survival strategy employed by bacteria in response to adverse environmental conditions.^24^ It has been reported that more than 100 species of bacterial species, including various *Bacillus* species, can enter the VBNC state under different environmental stresses, including exposure to nanomaterials.^25^ For instance, *P. aeruginosa* has been induced into the VBNC state under the stress of nanosilver, resulting in a significant decrease in culturable cells by six orders of magnitude at the concentration of 500 μg mL^-1^, while still maintaining relatively high metabolic activity.^12^ Therefore, it is reasonable to speculate that *B. tropicus* cells might enter the VBNC state under the treatment of BP nanosheets.

### 3.2 Calculation of the percentage of VBNC cells

To assess the proportion of VBNC cells, which are characterized by non-culturability and viability, we compared the number of viable cells to that of culturable cells.^13^ We quantified the amount of viable bacterial cells by measuring ATP concentration, as it serves as an indicator of metabolic activity.^16^ The measurement of ATP concentration disclosed moderate damage to *B. tropicus* caused by BP nanosheets, in contrast to the CFU method (Figure 2e). After treatment with 100 μg mL^-1^ of BP nanosheets, the energy metabolism of bacterial cells was significantly suppressed initially (at 4 h). However, the bacterial cells gradually adapted to the adverse environment and recovered their metabolic capacity over time. After 12 h, the ATP concentration accounted for approximately 60.93% of that in cells without BP nanosheets (Figure 2e). Under this circumstance, the number of viable cells, determined by the ATP concentration (60.93%), was 17.76 times higher than the number of culturable cells based on the CFU method (3.43%) (Figure 1b, c). This indicates that a large number of bacterial cells remained metabolically active despite their inability to form colonies. Subsequently, we calculated the percentage of *B. tropicus* cells in different states under the stress of BP nanosheets. At a concentration of 1 μg mL^-1^, there was minimal effect on bacterial growth activity. However, as the concentration increased, a greater number of bacterial cells entered the VBNC state. At the concentration of 100 μg mL^-1^, the culturable cells accounted for 4.05% of the total cells, while dead and VBNC cells accounted for 28.05% and 67.91%, respectively, compared to the control (Figure 2g).

Previous studies assessing the bacterial toxicity of BP nanosheets have typically relied on methods such as growth curve analysis, the CFU method, and morphological observation.^9-11, 22^ However, these methods often fail to detect the presence of bacteria in the VBNC state, leading to an oversight of the VBNC state in bacteria exposed to BP nanosheets. In contrast, our findings clearly demonstrated that there were more VBNC cells than culturable cells when exposed to BP nanosheets (Figure 2g). This highlights the limitations of conventional methods that solely rely on culturability and emphasizes the necessity of considering the metabolic activity when assessing the bacterial toxicity of BP nanosheets.

### 3.3 Transcriptome analysis

In order to gain a deeper insight into the mechanisms involved in the induction of the VBNC state by BP nanosheets, we conducted transcriptome sequencing. This analysis identified 406 differentially expressed genes (DEGs), most of which were significantly enriched into cellular metabolism, transmembrane transport, and cell respiration based on Kyoto Encyclopedia of Genes and Genomes (KEGG) and gene ontology (GO) enrichment analysis. Notably, most of these genes were down-regulated under BP nanosheet stress, indicating a suppressive effect of BP nanosheets on the metabolic processes and cell respiration in *B. tropicus.* Further details of the transcriptome sequencing analysis can be found in the Supporting Information 2.1 (Figure S2, Table S2, S3, S4). Therefore, subsequent analysis focused on these pathways to explore the molecular mechanisms underlying the induction of VBNC state by BP nanosheets.

### 3.4 Metabolic response to BP nanosheets

VBNC cells possess an extraordinary ability to maintain viability even with low metabolic activity.^26, 27^ When bacteria encounter harsh environmental conditions, they can enter a VBNC state, which allows them to sustain basal metabolic rate by down-regulating the expression of genes involved in central metabolism.^17^ Under BP nanosheet stress, a comprehensive analysis identified 72 pathways, with 69 showing up-regulation and 178 showing down-regulation of DEGs compared to the control group. Among these pathways, 51 were found to be related to metabolism, with 60 up-regulated and 113 down-regulated DEGs. Notably, the number of down-regulated DEGs in metabolic pathways was significantly higher than the up-regulated DEGs, indicating that BP nanosheets can suppress the metabolic activity of *B. tropicus* cells (Figure S2c, d).

One of the metabolic pathways that showed significant down-regulation under BP nanosheet stress was riboflavin metabolism (ko00740), with 4 out of 7 genes showing significant down-regulation (Table S5). This down-regulation could potentially hinder cellular respiration, as discussed in Section 3.5. In addition, valine, leucine, and isoleucine degradation (ko00280) displayed enrichment with 2 up-regulated and 4 down-regulated DEGs under BP nanosheet stress (Figure 3a, Figure S4, Table S5). Amino acids play a crucial role as building blocks for bacterial life, as they are necessary components of proteins involved in various cellular functions.^28^ Valine, leucine, and isoleucine are essential branched-chain amino acids, which can only be synthesized by microorganisms and make significant contributions to bacterial biomass.^28, 29^ The decreased synthesis of these branched-chain amino acids may, to some extent, hinder bacterial proliferation.

**Figure 3.**
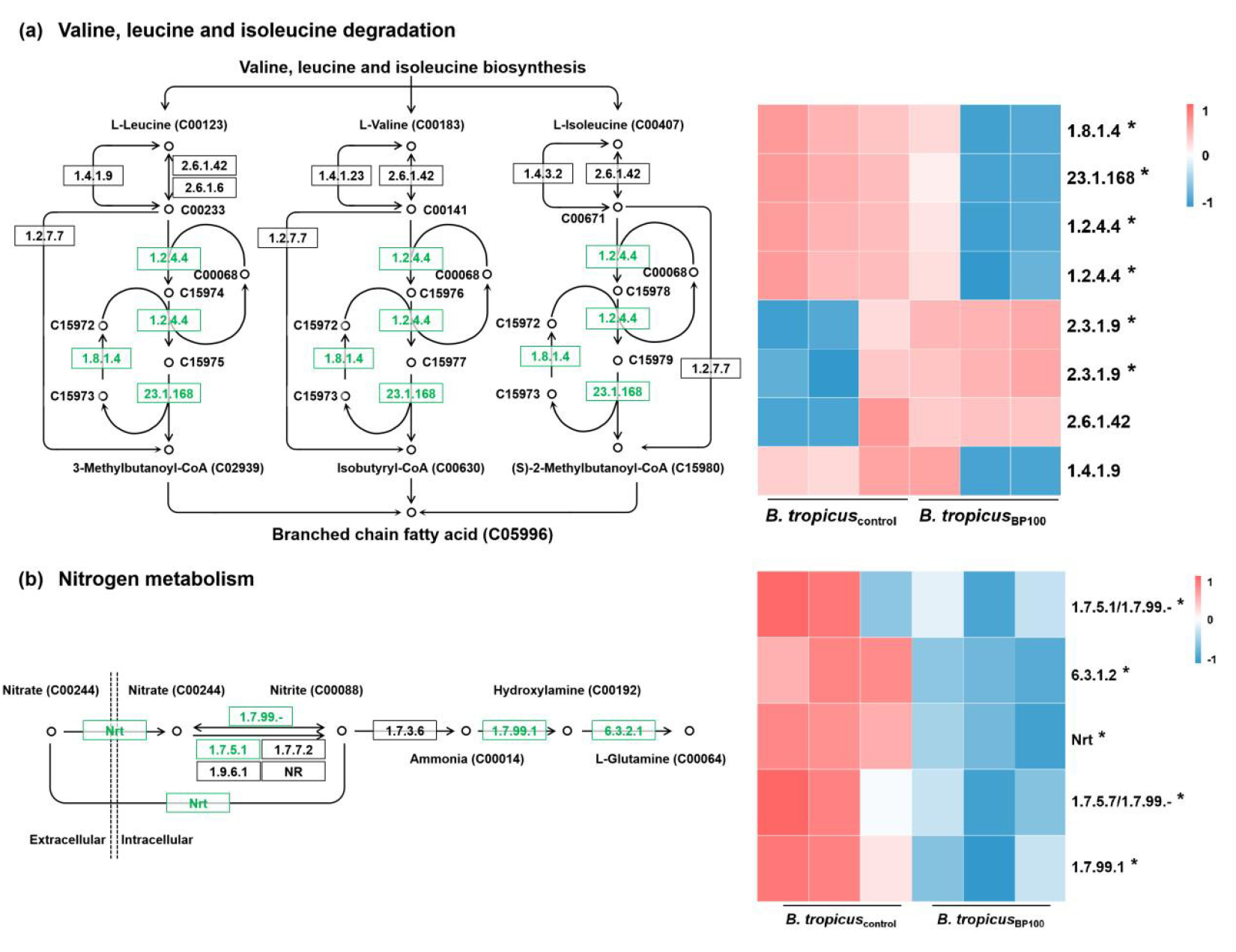
Effect of BP nanosheets on valine, leucine, and isoleucine degradation (a) and nitrogen metabolism (b). The heatmap represents the gene expression levels of key enzymes involved in these pathways. The circles and adjacent numbers represent the chemical compounds and their corresponding identifiers in the KEGG database, respectively. The boxes represent the enzymes responsible for catalyzing the respective steps. Genes with a green background indicate significant down-regulation compared to the control group.

In addition, nitrogen metabolism (ko00910) and propanoate metabolism (ko00640) were significantly enriched under BP nanosheet stress, with 5 and 8 down-regulated DEGs, respectively (Table S5). In the nitrogen metabolism pathway, the majority of genes were involved in the uptake and transformation of extracellular nitrogen (Figure 3b). Previous studies have highlighted that bacterial cells in the VBNC state exhibit a decreased rate of nitrogen utilization and maintain low nitrogen metabolic activity.^30^ Recently, Pan et al. found that bacterial cells down-regulated multiple pathways, including oxidative phosphorylation, arginine and proline metabolism, butanoate metabolism, and propanoate metabolism, in response to adverse environmental conditions.^31^ In our study, all of these pathways, expect for butanoate metabolism, were down-regulated, with 8 DEGs in oxidative phosphorylation, 3 DEGs in arginine and proline metabolism, and 8 DEGs in propanoate metabolism (Table S5). These results indicate that *B. tropicus* cells are likely to enter the VBNC state under BP nanosheet stress.

### 3.5 Cellular respiration and electron transfer under BP nanosheet stress

The down-regulation of riboflavin metabolism, as well as riboflavin synthesis and transport (GO:0006771, GO:0009231, and GO:0032218) under BP nanosheet stress suggested a decrease in riboflavin synthesis (Figure 4a, c, Table S4). Riboflavin is well-known for its crucial role in various biological processes. In bacteria, it acts as a key cofactor, supporting approximately 17% of enzymes that require cofactors.^32, 33^ Riboflavin serves as a precursor for flavin adenine dinucleotide (FAD) and flavin mononucleotide (FMN), which are crucial cofactors involved in several metabolic reactions.^34, 35^ In the electron transport chain, FMN acts as the first receptor, receiving electrons from NADH and transferring them to iron-sulfur clusters, which contain two-iron two-sulfur,^36^ four-iron four-sulfur,^37^ and three-iron four-sulfur.^35^ Ultimately, these electrons are transferred to bound quinone, a process that occurs in complex I.^35^ Therefore, any disruption in riboflavin metabolism and synthesis can affect the content and function of FMN, thereby impacting the electron transfer activity in the respiratory chain. In addition, under BP nanosheet stress, both four-iron four-sulfur cluster binding (GO:0051539, *p* = 0.029) and two-iron two-sulfur cluster binding (GO:0051537, *p* = 0.48) were down-regulated (Figure 4b, d). These findings indicate that BP nanosheets can inhibit electron transfer activity in the respiratory chain, potentially affecting riboflavin metabolism and the activity of iron-sulfur clusters.

**Figure 4.**
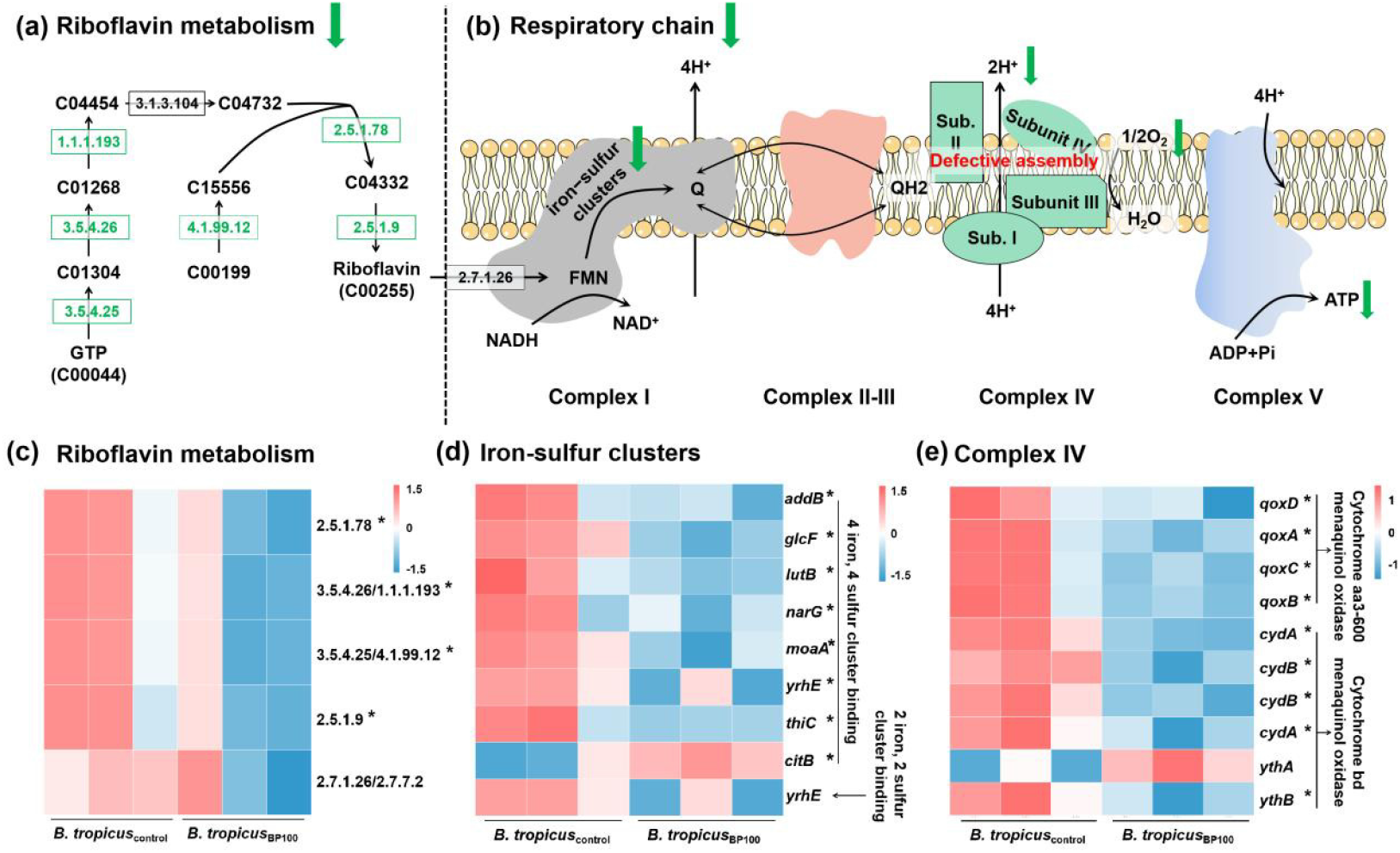
Riboflavin metabolism (a) and oxidative phosphorylation (b) in response to BP nanosheet stress. The heatmaps represent the gene expression levels related to riboflavin metabolism (c), iron-sulfur cluster (d), and complex IV (e). The numbers represent the identifiers of chemical compounds in the KEGG database. The boxes represent the enzymes responsible for catalyzing the respective steps. Genes with a green background indicate significant down-regulation compared to the control group.

BP nanosheets not only inhibited complex I in the respiratory chain, but also disrupted the assembly of complex III (Figure 4b). KEGG enrichment analysis revealed down-regulation of oxidative phosphorylation (ko00190) under BP nanosheet stress, with 8 significantly down-regulated DEGs identified (Table S5). These DEGs were identified as cytochrome *aa_3_*-600 menaquinol oxidase (4 DEGs) and cytochrome *bd* ubiquinol oxidase (4 DEGs). Bacterial oxidases are classified into four families (*o*, *d*, *a_1_*and *aa_3_*) based on the heme component they bind.^38^ In *Bacillus* species, two different terminal oxidases, *caa_3_*-605 and *aa_3_*-600, have been identified.^39^ Under BP nanosheet stress, the expression of genes related to *aa_3_*-600 was decreased (Figure 4e). This terminal oxidase acts as an electron donor and catalyzes the oxidation of quinol.^39^ The *aa_3_*-600 quinol oxidase is encoded by the *qox* operon, which includes four subunits (I, II, III, and IV) encoded by the genes *qoxB*, *qoxA*, *qoxC*, and *qoxD*, respectively.^40, 41^ Villani et al. have demonstrated the importance of *qoxC* and *qoxD* in the proper assembly and proton pumping of *aa_3_*-600 quinol oxidase.^40^ In *B. tropicus*_BP100_, the expression of *qoxC* and *qoxD* was found to be 3.87 and 3.10 times lower, respectively, compared to *B. tropicus*_control_ (Table S6). Under BP nanosheet stress, defective assembly and decreased proton pumping of *aa_3_*-600 quinone oxidase may occur. However, *Bacillus* species also have an alternative branch of the respiratory chain involving a quinol oxidase, known as the *bd* terminal oxidase, which catalyzes proton transfer and contributes to cell respiration.^42^ Under BP nanosheet stress, the expression of *cydA* and *cydB,* which encode the cytochrome *bd* terminal oxidase,^42, 43^ was significantly down-regulated (Table S6). The inhibition of these genes may affect the expression cytochrome *bd* and ultimately lead to a decrease in bacterial cell respiration.^42^

Under aerobic conditions, bacterial cells can produce ATP through the respiratory chain.^44^ In our study, GO enrichment analysis revealed significant suppression of several terms, including electron transfer activity (GO:0009055, *p* = 0.0022), ATP synthesis coupled with electron transport (GO:0042773, *p* = 0.0017), respiratory electron transport chain (GO:0022904, *p* = 0.0038), and oxidative phosphorylation (GO:0006119, *p* = 0.0032). These findings indicate that BP nanosheets inhibited electron transfer in the respiratory chain. During the electron transfer process in the respiratory chain, from complex I to IV, ATP synthesis is driven by an electrochemical gradient, providing the necessary energy for ATP synthesis .^35^

To investigate the potential inhibition of ATP production in *B. tropicus* as a result of suppressed respiratory chain activity, we conducted a cell metabolism test using the SeaHorse XF24 analyzer. This platform allows for the real-time measurement of oxygen consumption rate (ORC) at a picomole resolution, which serves as an indicator of cellular respiration.^44^ After treatment of *B. tropicus* with BP nanosheets, we observed a decrease in cellular respiration compared to the control group (Figure S5). When bacterial cells enter the VBNC state, they may undergo metabolic and physiologic changes, such as a reduction in cellular respiration, as a means to maintain cellular activity.^45^ This adaptation could potentially enable *B. tropicus* to survive under BP nanosheet stress.

### 3.6 Resuscitation of VBNC bacteria

Although VBNC bacteria lose the ability to grow on normal culture medium, it is important to note that this does not imply their death. Resuscitation, which refers to the recovery of VBNC cells, has been recognized as an important characteristic.^26^ When the environmental stress is removed and the cells are cultured under suitable conditions, bacterial cells can be revived.^14^ In our study, transmission electron microscopy (TEM) images revealed significant protein aggregation in *B. tropicus* cells (Figure 5a, d). However, after 12 hours of treatment with BP nanosheets, most *B. tropicus*_BP100_ cells exhibited reduced protein aggregation with more voids (Figure 5b,e). This observation aligns with previous studies that have shown a slowdown nutrient absorption, a reduction in macromolecular synthesis and metabolism, as well as a decrease in cytoplasmic and total protein concentrations during the transition from the normal state to the VBNC state under environmental stress.^14, 46^ After removing the BP nanosheet stress and culturing for an additional 12 h, most resuscitated cells demonstrated recovered protein synthesis and aggregation (Figure 5c, f). Although the protein density in the resuscitated cells was not as dense as in the control cells, our data provided preliminary evidence that BP nanosheets do not kill the gram-positive bacterium. However, they have the potential to induce bacteria to enter the VBNC state.

**Figure 5.**
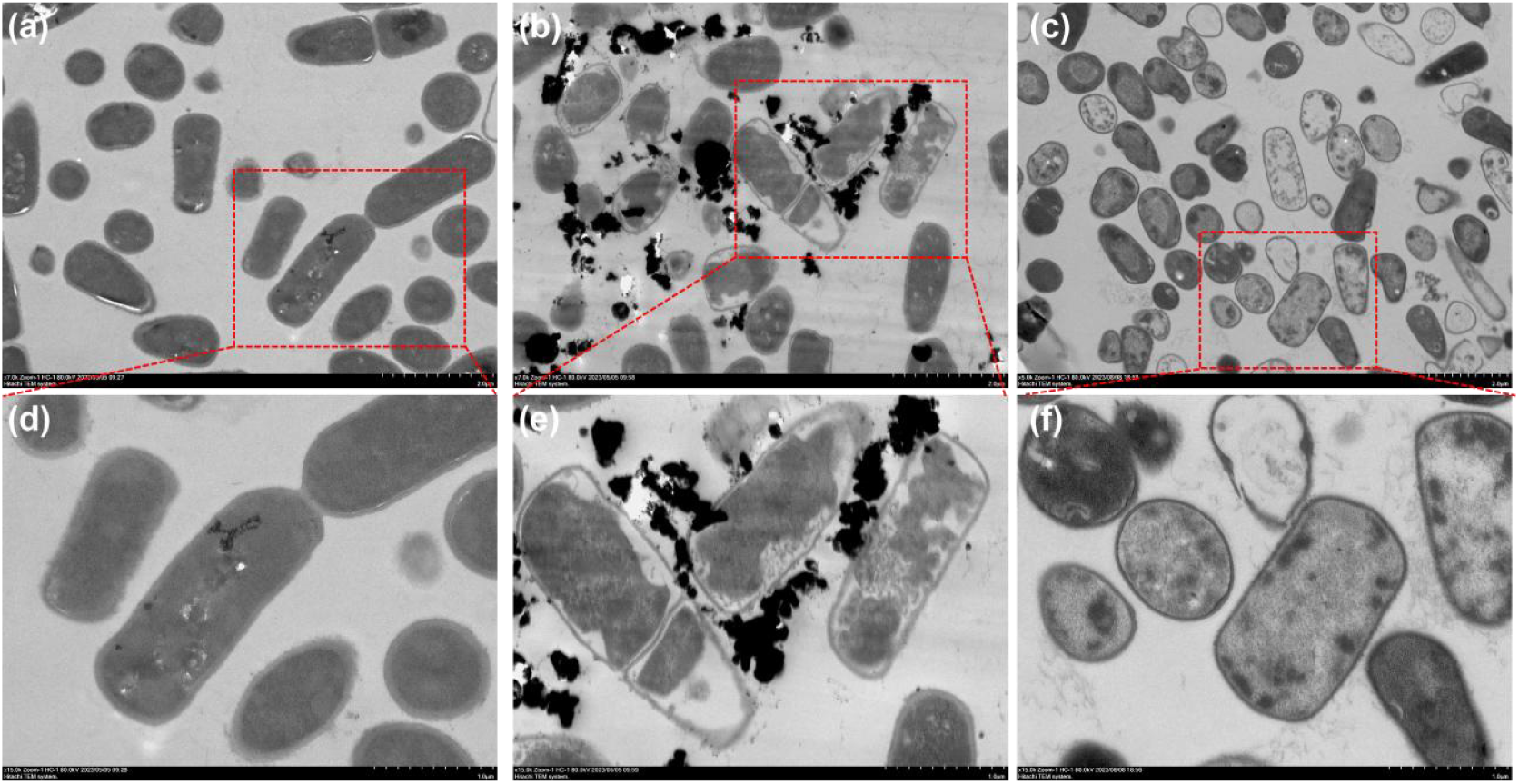
TEM images of *B. tropicus* without any treatment (a, d), after treatment with BP nanosheets for 12 h (b, e), and following the removal of BP nanosheet stress (c, f).

### 3.7 Antibiotic susceptibility test

Due to the lower metabolic activity and basal respiration of bacterial cells in the VBNC state, they may exhibit greater antibiotic resistance compared to culturable cells. This is primarily because the reduced basal respiration prevents antibiotics from inducing tricarboxylic acid cycle (TCA) activity, thereby avoiding metabolic toxicity and minimizing lethality.^47^ In our study, BP nanosheets were found to down-regulate several GO terms related to cellular metabolism, including respiratory electron transport chain (GO:0022904), oxidative phosphorylation (GO:0006119), drug transport (GO:0015893), drug transmembrane transport (GO:0006855), and response to drug (GO:0042493) (Table S4). Combined with the lower metabolic activity, we speculate that BP nanosheets may increase the tolerance of *B. tropicus* to antibiotics.

To confirm this conjecture, we pretreated *B. tropicus* cells with BP nanosheets (100 μg mL^-1^) and then examined their tolerance to exogenous antibiotic treatment. We observed concentration-dependent growth inhibition for both *B. tropicus*_control_ and *B. tropicus*_BP100_. After kanamycin stress for 30 min, a slight decrease in bacterial activity was observed in *B. tropicus*_control_ compared to *B. tropicus*_BP100_, as determined by ATP measurements at doses of 25 and 50 μg mL^-1^. However, the decrease in bacterial viability was more pronounced in *B. tropicus*_control_ (40.22% ± 2.62% and 34.26% ± 2.91%), compared to *B. tropicus*_BP100_ (45.98% ± 2.31% and 40.90% ± 3.07%) after 60 min (Figure S6a). Moreover, through OCR measurements, we observed that the decrease in bacterial respiration in *B. tropicus*_control_ was more significant than in *B. tropicus*_BP100_ following kanamycin treatment (Figure S6b). These results collectively indicate that BP nanosheets reduced the susceptibility of *B. tropicus* to kanamycin. Previous studies have demonstrated that VBNC cells possess a robust capability to withstand antibiotic treatments.^48–50^ In a mouse model of urinary tract infection, Bryan Rivers and Todd R. Steck observed VBNC cells persisting after antibiotic treatment. Notably, these cells were capable of resuscitation within the host once the antibiotic treatment ceased,^48^ demonstrating the resilience of VBNC cells under antibiotic stress. Advances in transcriptome sequencing technology enabled researchers to discover that *Bacillus subtilis* upregulated certain genes involved in proline uptake and catabolism in response to kanamycin stress.^51^ In the current research, it was noted that three genes associated with arginine and proline metabolism were downregulated after exposure to BP nanosheets (Table S5), which could potentially enhance the tolerance to antibiotics. However, the precise implications of this finding need further investigation. In addition, since bacterial metabolism in the presence of antibiotics involves numerous, complex, and coordinated biomolecular networks,^47^ it becomes challenging to predict which antibiotics can effectively combat bacteria under BP nanosheet stress. Nevertheless, our study provides a preliminary understanding of the need to considerate the changes in the gene regulatory network that may lead bacteria to enter the VBNC state and decrease their susceptibility to antibiotics.

## 4. Conclusion

In summary, this study aimed to investigate the response of *B. tropicus* to BP nanosheets and reveal the underlying molecular mechanisms at the transcriptional level. Our observations indicated that, rather than being killed, the majority of bacterial cells entered a VBNC state when exposed to BP nanosheets. The nanosheets induced respiratory depression and a decrease in cellular metabolism, as indicated by the down-regulation of pathways related to the respiratory chain and cellular metabolism. These findings not only deepen our understanding of the biological effects of BP nanosheets on bacteria and provide insights for their ecological risk assessment, but also highlight the significance of utilizing appropriate approaches in evaluating the bacterial toxicity of nanomaterials.

## Acknowledgements

This work was supported by the National Natural Science Foundation of China (No. 42022056, 42192574, 42277423, 42077394, 22176196), GDAS’ Project of Science and Technology Development (2022GDASZH-2022 010105 and 2020GDASYL-20200101002), and Guangdong Foundation for Program of Science and Technology Research (No. 2020B1212060048).

